# Basal forebrain-derived acetylcholine encodes valence-free reinforcement prediction error

**DOI:** 10.1101/2020.02.17.953141

**Authors:** JF Sturgill, P Hegedus, SJ Li, Q Chevy, A Siebels, M Jing, Y Li, B Hangya, A Kepecs

## Abstract

Acetylcholine (Ach) is released by the cholinergic basal forebrain (CBF) throughout the cortical mantle and is implicated in behavioral functions ranging from arousal to attention to learning. Yet what signal ACh provides to cortex remains unresolved, hindering our understanding of its functional roles. Here we demonstrate that the CBF signals unsigned reinforcement prediction error, in contrast to dopamine (DA) neurons that encode reward prediction error. We show that both CBF neuronal activity and acetylcholine (ACh) release at cortical targets signal reinforcement delivery, acquire responses to predictive stimuli and show diminished responses to expected outcomes, hallmarks of a prediction error. To compare ACh with DA, we simultaneously monitored the activity of both neuromodulators during a serial reversal learning task. ACh tracked learning as swiftly as DA during acquisition but lagged slightly during extinction, suggesting that these neuromodulators play complementary roles in reinforcement as their patterns of innervation, cellular targets, and signaling mechanisms are themselves complementary. Through retrograde viral tracing we show that the cholinergic and dopaminergic systems engage overlapping upstream circuits, accounting for their coordination during learning. This predictive and valence-free signal explains how ACh can proactively and retroactively improve the processing of behaviorally important stimuli, be they good or bad.

## Introduction

The cholinergic basal forebrain (CBF) is positioned to profoundly influence brain function through its projections that extend throughout pallium-derived structures including the cortex, amygdala and hippocampus (*1, 2*). Pharmacological, lesion, and optogenetic evidence implicates the CBF in an exceedingly diverse array of processes ranging from sleep, arousal, movement, attention, learning, memory to cognitive decline(*3–8*). In particular, forms of perceptual, skill, and emotional learning require cholinergic modulation of extra-striatal structures(*9–11*).

Like ACh, DA has been implicated in a diverse set of seemingly incompatible functions, including movement, pleasure and learning(*12*). Recordings from midbrain dopamine neurons provided a critical breakthrough, revealing that dopamine neurons obey a simple computational principle, signaling rewarding prediction error (RPE), i.e. they fire when reward expectations are surpassed(*12*). In contrast to DA, we know comparatively little about what signal ACh encodes during behavior, as only recently have direct recordings of CBF activity become technically feasible(*6*–*8, 13*–*15*). Previous theoretical work suggested that ACh and other neuromodulators play distinct and independent computational roles in reinforcement learning(*16*). In particular, ACh was proposed to track uncertainty and to control learning rate(*17, 18*). Yet evidence for a computational principle that might help reconcile the CBF’s diverse functions is lacking.

Once thought to gradually and diffusely promote arousal and attentive states, recent evidence instead shows that phasic, temporally-precise CBF activity is triggered by rewards and salient stimuli (*13, 19*). Inspired by these findings, we considered the possibility that the CBF provides reinforcement predictions to cortical areas analogously to dopaminergic reward prediction error signals transmitted to the ventral striatum(*12, 20, 21*). However, if the CBF were to convey a signed, value-based signal it would imply the existence of cortical mechanisms for interpreting and distinguishing positive and negative neuromodulator fluctuations. Cortex contains no obvious counterpart to the segregated type 1 and 2 DA receptor-expressing populations of spiny projection neurons shown to serve this function within striatum(*22*). By contrast, a valence-free reinforcement prediction error would provide an alerting signal appropriate to the modular and functionally heterogeneous organization of cortex yet necessitate separate pathways for distinguishing reward and punishment.

## Results

Therefore we resolved to determine whether the CBF encodes prediction errors and, if so, to ascertain their computational form. Mice were trained to associate odor cues with reinforcement in a Pavlovian task and odor cue-evoked licking was used to index reward expectation(*20*). We then examined whether prediction errors were encoded in the spiking of identified cholinergic neurons within the horizontal limb of the diagonal band (HDB) using optogenetic tagging (*13*). To impart different expected values to distinct odorants, we varied the outcome probabilities for reward (H20), punishment (air puff), or omission. Demonstrating that CBF neurons encode reinforcement predictions, odor cues elicited phasic firing in proportion to expected value (Fig. 1A). Consistent with previous results, individual CBF neurons responded to reward delivery with brief bursts of spikes(*13*). Moreover, reward-evoked spike rates were diminished when preceded by the high value cue relative to the low value cue (Fig. 1A).

**Figure 1.**
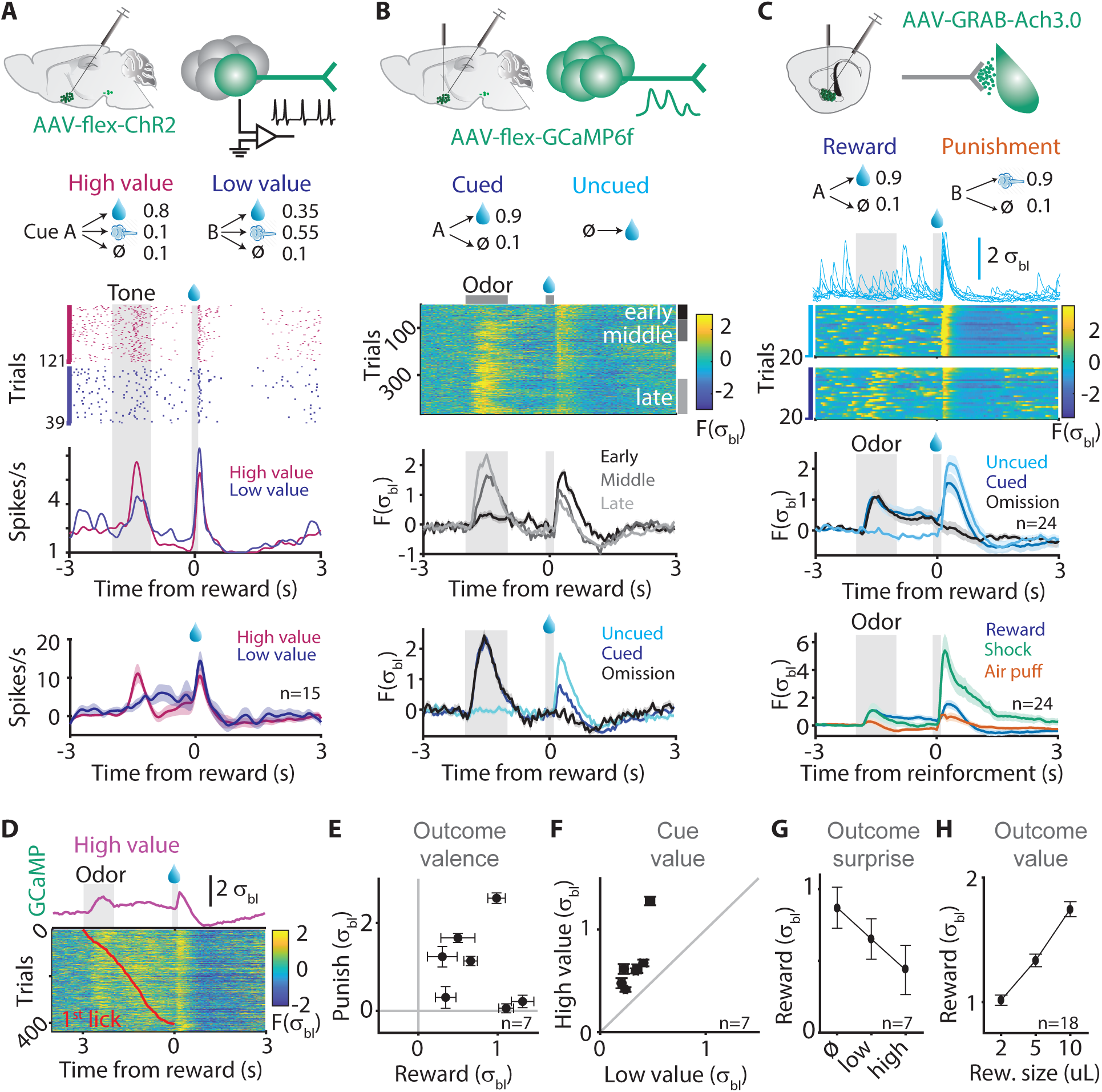
Basal forebrain cholinergic neurons encode and transmit unsigned reinforcment prediction errors. **A** Top, schematics for optogenetic tagging and recording of CBF neurons within the HDB and for probabilistic outcome task with cue and outcome probabilities indicated. Middle, example spike raster and histogram for rewarded high- and low-value odor trials. Evident are phasic cue- and reward-evoked spike responses. Bottom, grand average spike histogram. **B** Top, fiber photometry recording from cre+ neurons virally transduced with GCaMP6f in HDB. Middle, photometry raster plot of cued, reward trials showing gradual acquisition of CBF responses to reward predictive cue. Color bars indicate representative early, middle, and late training sessions for accompanying averages. Bottom, at steady state, reward predictive cues trigger CBF activation and lead to diminished responses to reward delivery, as shown from averages for a representative mouse. **C** Top, schematics for monitoring ACh fluctuations at axonal targets within the BLA and for cued reward and punishment task. Middle, reward evokes transient increases in extracellular ACh (traces at top) that are reliable across trials and diminished by reward predictive cue (rasters at bottom). Grand averages show diminished responses to expected relative to unexpected reward (** p<0.005, cued vs. uncued). **D-G** Responses of HDB cre+ neurons expressing GCaMP6f recorded during probabilistic outcome task. **D** Representative photometry raster plot for high value, reward trials sorted by response latency (red line) with corresponding average shown above. Onset of CBF activity precedes anticipatory licking. **E** Positive responses to reward (water) and punishment (air puff). **F** CBF responses to high value odor cue are consistently greater. **G** Reward-evoked responses are inversely proportional to reward expectation. **H** Reward-triggered ACh release in the BLA is proportional to reward size. Error bars, S.E.M.

To efficiently record CBF activity at the population level and track its evolution during learning, we used fiber photometry to record bulk intracellular Ca^2+^ from cre+ neurons expressing GCaMP6f using ChAT-cre mice (*23*). CBF neurons initially responded selectively to reward delivery and not to novel odorants but acquired responses to reward-predictive cues with learning (Fig. 1B, fig. S1). While correlated with licking, alignment to lick onset revealed that cue responses preceded licking and were better explained by stimulus onset (fig. S2). Corroborating our electrophysiological findings, population Ca^2+^ responses to reward appeared diminished for predicted relative to surprising rewards (Fig. 1B).

Direct recordings of cholinergic transmission provided further confirmation that the CBF signals prediction errors. Such recordings were crucial because cholinergic transmission might conceivably be gated locally via presynaptic modulation, decoupling spiking activity from synaptic release, and the quantitative relationships between CBF Ca^2+^, spike rate, and neurotransmitter release are not well understood(*24, 25*). To measure CBF-evoked extracellular ACh release we introduced the improved fluorescent ACh sensor, ACh3.0(*26*), within principal cells of the basolateral amygdala (BLA), a major axonal target of the CBF. In the absence of a stimulus, brief fluorescence transients stochastically appeared within single trials, consistent with sparse and/or synchronous activation of cholinergic terminals within range of the ∼200um radius recording volume (Fig. 1C)(*27*). By contrast, high baseline firing and/or spatial pooling of extracellular ACh would be expected to obscure such sparse activity through summation. Reward triggered rapid and temporally aligned transients followed by a pause, marked by the absence of spontaneous transients immediately following reward delivery as well as a drop and gradual recovery of fluorescence (Fig. 1C). Cue delivery elicited phasic increases in extracellular ACh and responses to expected relative to unexpected reward were clearly and significantly diminished (p<0.005; n=24 fibers; Fig. 1C). Thus, we confirmed that CBF activity is faithfully conveyed to axonal targets as reflected by terminal ACh release. Taken together, these data show that CBF activity bears the cardinal features of an unsigned prediction error: reinforcement response, prediction, and surprise.

Additional experiments resolved that the CBF signals an unsigned, valence-free prediction error that quantitatively reflects reinforcement magnitude. First, punishment-predictive cues as well as punishment delivery consistently triggered ACh release in the BLA. ACh release was stronger for more intense (tail shock) relative to more mild (air puff) reinforcement stimuli (Fig. 1C) and was fast and precise (fig S3)(*13*). Similarly, CBF neurons in the HDB were phasically activated by both punishment and reward (p<0.005; n=7 mice; Fig. 1E)(*13*). Second, CBF activity quantitatively reflected expected value. In the probabilistic outcome task, cue-evoked Ca^2+^ responses were consistently greater for the high value vs. low value odor (p<0.005; n=7 mice; Fig. 1F). In addition, the reward-triggered step increase in fluorescence was diminished in proportion to cue value (p<0.05 low vs high; n=7 mice; Fig. 1G). Furthermore, reward-evoked ACh transients in the BLA were monotonically related to reward size (p<0.05; n=18 fibers; Fig. 1H).

Are cholinergic prediction error signals consistently delivered across cortical regions and are they coordinated in time? The local variation of cholinergic signals at axonal targets remains controversial and poorly characterized due to technical limitations (*13, 19, 28*). Using the ACh3.0 sensor to simultaneously record ACh fluctuations from different locations, we uncovered a remarkable coordination of Ach release. Even across hemispheres, extracellular ACh in the BLA was temporally aligned and highly coherent (Fig. 2A-C). Encouraged by this observation, we investigated whether CBF-derived prediction error signals were consistently broadcast across brain areas by delivering predictive cues and reinforcers (water reward, air puff, tail shock) that varied in sign, magnitude, and identity (Fig. 2D). To account for differences in viral expression or fiber placement, we normalized all responses to uncued reward and quantified the covariance of their means (signal correlations) as well as of their residuals (noise correlations) (Fig. 2D). Relative to unexpected reward, mean responses to other stimuli tended to scale proportionately (Fig. 2E). Furthermore, mean response magnitudes were positively correlated, more so within than across animals (mean R=0.78; n=11 mice; Fig. 2F). Yet in a subset of cases, we observed significant biases across hemispheres. Mean responses to aversive stimuli, in particular, could deviate from reward (Fig. 2E). Thus the CBF signal is broadly consistent yet exhibits some unaccounted for heterogeneity.

**Figure 2.**
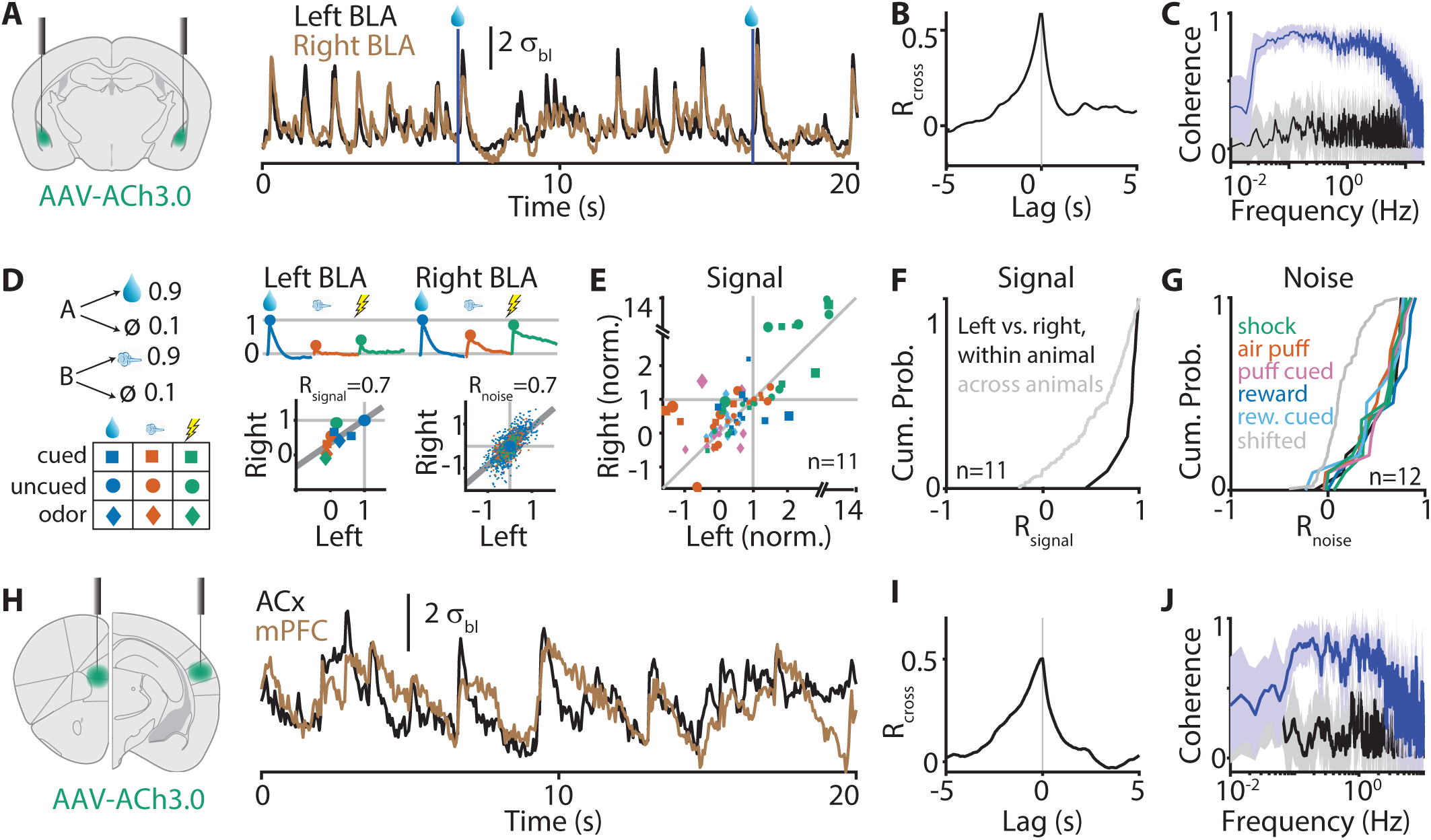
The CBF provides consistent and coherent reinforcement signals to axonal targets. **A** Recording schematic (left) and example traces (right) for simultaneous measurements of extracellular ACh from locations within the left and right BLA. **B**,**C** Strongly correlated and coherent ACh3.0 fluorescence responses recorded across hemispheres. **D** Left, behavioral task schematic. Top, mean cue and outcome signals normalized to uncued reward shown for an example mouse. Bottom, positive signal (left) and noise (right) correlations for same. **E** Scatter plot shows mean normalized responses pooled across mice. Unity line indicates equal response magnitudes relative to uncued reward. Marker size proportional to effect size (D’) of bias or distance from unity line. **F** Cumulative probability across mice of signal correlations between the left and right BLA. Grey line shows null distribution of correlations across animals where left and right recordings sites were mismatched by permutation. **G L**argely positive cumulative distributions of noise correlations separated by reinforcement condition. **H-J** As for A-C but for simultaneous recordings from ACx and mPFC.

Moreover, CBF signals are coordinated in time as well as space as trial to trial variation in stimulus responses was highly correlated. Even across hemispheres, we recorded remarkably high noise correlations that accounted for a substantial portion of the total variance (38% median; n=12 mice) of reinforcement-evoked fluorescence transients recorded using the ACh3.0 sensor (Fig. 2E). These correlations were only weakly driven by gradual drift or session-wide trends (e.g. satiety or photobleaching) as shifting the data set by one trial presentation markedly reduced the total variance explained (1.8% median; n=12 mice). Such strong noise correlations indicate that CBF neurons share common upstream inputs and/or reciprocal connections. Correlated ACh release implies that its effects on cortical desynchronization and modulation of sensory responses are tightly controlled and suggests that ACh significantly contributes to the spatiotemporal coordination of brain activity(*10, 29*). Importantly, tightly coordinated ACh release is not unique to the BLA or to homologous brain regions *per se*. Simultaneous recordings from medial prefrontal cortex (mPFC) and primary auditory cortex (ACx) revealed a comparable synchronization evident as a peaked cross correlogram and a broad extent of significantly positive coherence magnitude (Fig. 2H-J).

That the CBF encodes a reinforcement prediction error suggests a role in reinforcement learning, yet leaves unclear whether CBF signals track or lag changes in conditioned behavior. To quantify CBF plasticity relative to the dopaminergic VTA while controlling for variable learning rates across behavioral sessions, we took advantage of the anatomical separation of CBF and VTA DA neurons to record both populations simultaneously in ChAT(cre/+);DAT(cre/+) mice. Initially, we trained mice in an odor-cued serial reversal task featuring positive (CS+) and negative (CS-) value reinforcement contingencies (valence task) and performed repeated reversals across days (Fig. 3A). Mice quickly adapted to new contingencies, requiring 11 trials for acquisition and 19 trials for extinction on average to reach 80% of asymptotic performance. Unlike VTA DA, initial CBF cue responses to the aversive CS-were rectified at zero, but both systems rapidly adapted following the reversal point, increasing their similarity (Fig. 3C, top). By contrast, extinction of VTA DA cue responses appeared more sudden compared to those of the CBF, supported by a window of divergence following reversal of the former CS+ cue (Fig. 3C, bottom). To account for the possibility that aversive stimuli might differently engage neural circuits mediating learning and produce distinct neuromodulatory dynamics, we trained another cohort of mice on a separate task involving alternate valuation/ devaluation of odor cues (value task) and attained broadly similar results (fig. S4).

**Figure 3.**
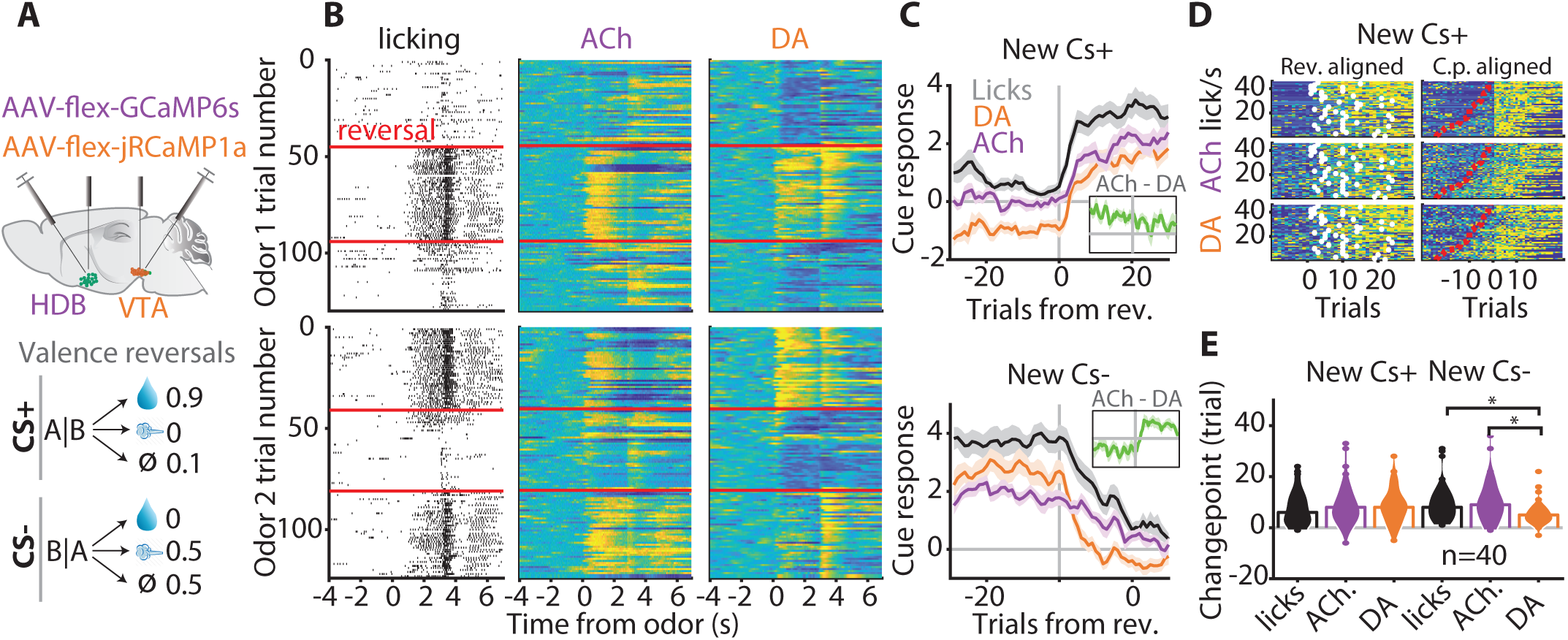
Cholinergic and dopaminergic predictions adapt with highly similar dynamics during learning. **A** Recording schematic for simultaneous recordings (top) and outcome contingency table for serial valence reversal task (bottom). **B** Example raster plots for licking and ACh and DA neuron activity for odor 1 (top) and 2 (bottom), red lines indicate reversal trials. **C** Average cue responses for newly (new Cs+, top) and formerly (new Cs-, bottom) rewarded odors show that ACh dynamics are qualitatively similar to DA. Inset shows difference between ACh and DA signals, note step increase for new Cs-trials. **D** Quantification of learning speed per reversal using changepoint detection. Left, responses aligned to reversal point with behavioral changepoint calculated from cue-evoked licking indicated by white markers. Right, alignment by changepoint with trials sorted by the same and reversals shown by red line. **E** Changepoint distributions for licks, ACh and DA. Distributions were statistically indistinguishable during acquisition whereas median DA changepoints occurred significantly earlier compared to licks and ACh during extinction.

To estimate the learning latency for each reversal while imposing minimal constraints, we applied changepoint detection to the cue-evoked anticipatory licking and fluorescence responses (Fig 3D, S5). Alignment of neural responses to the behavioral changepoint for each reversal revealed that both systems adapted indistinguishly across a 10 trial window during acquisition, but the CBF lagged VTA DA during extinction. Indeed, the average changepoint trials for licking, CBF, and DA activity were statistically indistinguishable during acquisition (N.S.; n=36 reversals; Fig. 3E) whereas ACh and licking significantly lagged DA during extinction (p<0.005; n=40 reversals; Fig. 3E). Thus, CBF predictions about reinforcement adapt sufficiently rapidly to participate in value updating and/or modification of conditioned behavior.

## Discussion

Contrary to preexisting notions of independent neuromodulatory roles, our data demonstrate that the CBF and dopaminergic VTA provide computationally related signals. The coordination of these signals across brain areas must therefore be crucial for shared behavioral functions such as learning. However, very little is known about how different neuromodulators are coordinated in space and time due to the technical challenge of recording multiple, identified cell types simultaneously across different brain areas. Motivating us to address this question, we observed that in reversal experiments, CBF and VTA DA responses not only encoded related prediction errors on average, but were further correlated on a trial-to-trial basis (Fig. 4A,B). Thus, using dual fiber photometry, we directly compared externally- and internally-generated patterns of CBF and VTA DA activity by delivering unpredictable rewards in an otherwise unstructured task environment. Surprisingly, we found the degree of coordination of reinforcement-driven and spontaneous activity to be qualitatively similar. Both neuromodulators closely tracked whisking and brief fluctuations in arousal whereas CBF activity appeared more closely associated with extended bouts of locomotion (Fig. 4C)(*6, 7*). Overall, their activity was strongly coherent across time scales of seconds to tens of seconds. Using time-resolved coherence estimates, we found that coherence was not solely driven by reinforcement, as it was maintained long after reward delivery (p<0.05; n=6 mice; Fig 4E,F).

**Figure 4.**
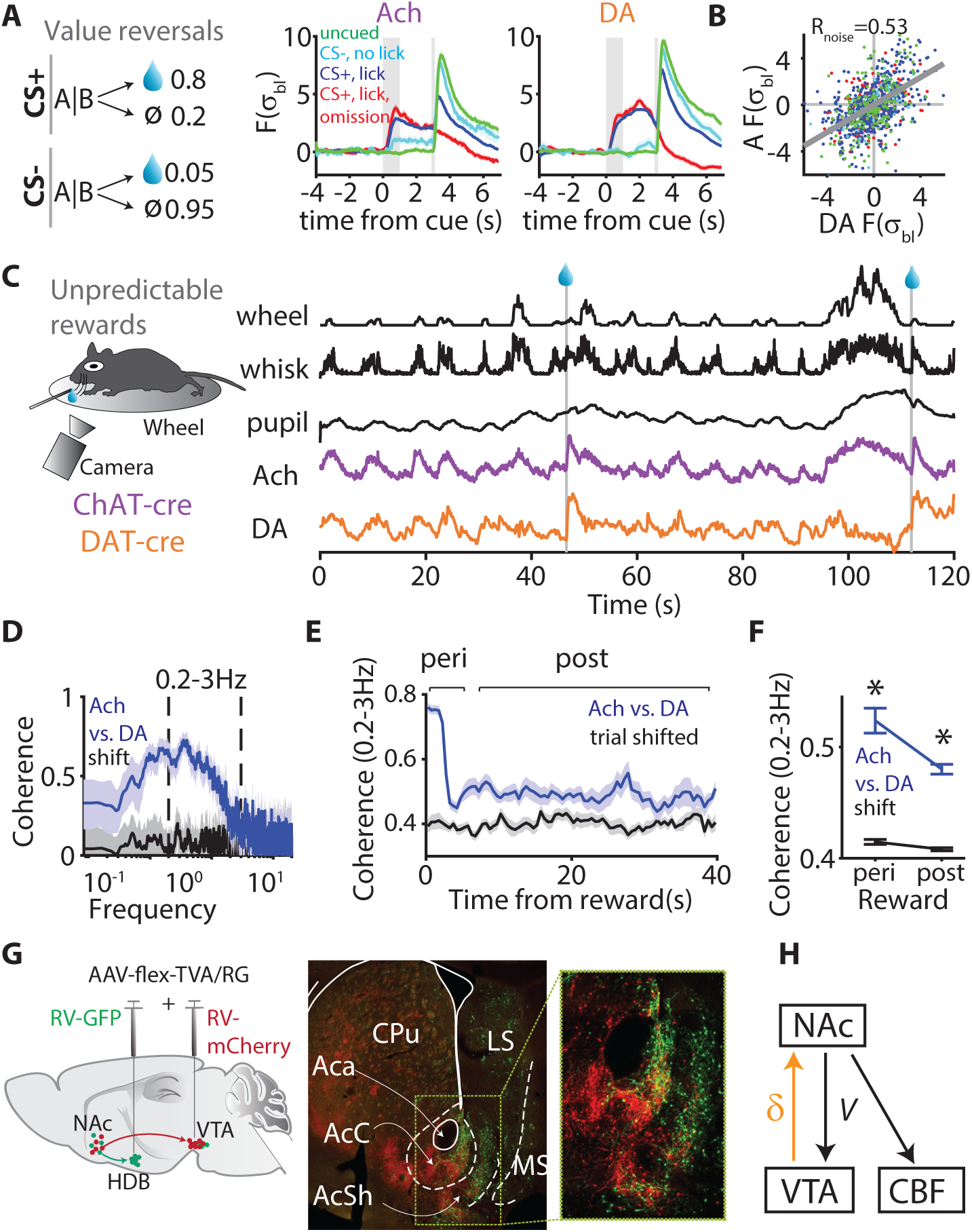
Computational, dynamical, and anatomical similarity of the cholinergic and dopaminergic neuromodulatory systems. **A** Average responses of simultaneously recorded CBF and VTA DA neurons encode reinforcement prediction errors across serial value reversals. **B** Scatter plot of mean-centered response magnitudes, linear fit, and noise correlation for same mouse as in panel A. **C** Random rewards task and example traces illustrating covarying CBF and VTA DA activity and correlations with spontaneous behavior. **D** Spectral coherence magnitude for CBF and VTA DA activity during spontaneous behavior **E**,**F** Time-resolved coherence estimates aligned to reward delivery show that CBF and VTA DA activity are strongly coupled by reinforcement (peri-reward) but still significantly coupled in the absence of external stimuli (post-reward). **G** Retrograde tracing reveals that both CBF and VTA DA neurons receive prominent input from partially overlapping regions of the NAc core and shell. **H** Circuit model proposing that NAc input provides a value signal responsible for related VTA DA and CBF reinforcement prediction errors.

To address what circuit mechanisms account for the coordination of CBF and VTA DA neuron activity we directly compared their sources of synaptic input via retrograde, transynaptic tracing with pseudotyped rabies virus (RV)(*30*). CBF neurons posterior to the nucleus accumbens (NAc) in the anterior nucleus basalis and VTA DA neurons were infected with AAV-flex-TVA and AAV-flex-RG viruses in ChAT-(cre/+);DAT-(cre+) mice, rendering them competent for PRV infection and retrograde, transynaptic labelling. Subsequent infection with GFP- or mCherry-encoding RV in the CBF and VTA, respectively, labelled populations of neurons distal to the injection site in both distinct and overlapping regions, including the prefrontal cortex, NAc, and lateral hypothalamus, consistent with previous findings (*31–34*). In particular, both neuromodulatory systems received input from overlapping neuronal populations within the NAc, with inputs to the CBF being biased towards medial regions of the NAc core and shell. Thus, overlapping and distributed sets of inputs likely provide information about value to CBF and VTA DA neurons, for the computation of prediction errors (*35*).

Here we clarify what signal CBF neurons convey to pallial targets, showing that CBF activity encodes reinforcement prediction errors. However, unlike canonical dopaminergic RPE, the CBF prediction error is unsigned and therefore valence-free. Consistent with an unsigned prediction error, recent work has demonstrated that CBF neurons projecting to the auditory cortex respond to auditory tones associated with foot shock(*14*). Unsigned prediction errors feature prominently in attentional learning theories where they effectively control the learning rate for associating cues with outcomes(*36, 37*). Indeed, manipulations of CBF activity has been demonstrated to affect performance in learning tasks(*9–11*). Roles for unsigned prediction errors are not exclusive to learning, however, they have also been proposed to elicit attention(*38*). CBF activity has been shown to correlate with both movement and arousal, properties that are notably shared to varying degrees with the dopamine system (*6*–*8, 39, 40*). While movement and arousal are inextricably tied to learning in a Pavlovian setting, our data additionally reveal a rapid, reliable and phasic CBF activation by salient sensory cues.

We provide new insights into the spatiotemporal organization of CBF neuromodulation. First, it is remarkably fast. Extending previous electrochemical measurements, we show that the temporal resolution of stimulus-evoked acetylcholine transients approaches 100ms (a possible underestimate given exogenous expression ACh3.0 reporter)(fig. S3)(*19*). Second, it is largely consistent in space and time. Trial by trial fluctuations in acetylcholine release were highly correlated, even across hemispheres, which may reflect an important role for acetylcholine in coordinating activity between spatially segregated brain regions. Furthermore, mean reinforcement responses were correlated across different target locations within an animal, much more so than across animals (Fig. 2D). Thus, inbred mice respond heterogeneously to reward and punishment and CBF activity reflects these inter-individual differences. Still, while responses to most stimuli in most animals were proportional, we observed sizeable biases in a subset of cases, e.g. punishment responses could be consistently stronger than reward responses at one recording site relative to the other. Such differences suggest a mosaic organization in which a small number of axons contribute to local acetylcholine sensed by individual neurons (given similar spatial scales for photometry recordings and dendritic arbors) and may reflect tuning differences in CBF neurons themselves or presynaptic modulation of cholinergic terminals(*27, 28*).

Our data supports parallel and coordinated roles in learning for ACh and DA, enabled by a distributed circuit for computing prediction errors. Both neuromodulatory systems adapted with highly similar dynamics during learning. However, CBF cue responses extinguished more slowly compared to VTA DA during serial value or valence reversals. This difference may highlight a selective role for ACh in extinction learning, consistent with increased perseverative responding following manipulations of CBF activity and analogous to the serotonergic system (*41–43*). Remarkably, we found that fluctuations in CBF and VTA DA activity as well as ACh release within the BLA were correlated. These findings support recent evidence for a highly distributed and redundant circuit for computing dopaminergic RPE(*35*), which may involve indirect callosal or subcortical commissural pathways as all of our simultaneous recordings were performed across hemispheres. Based upon shared connectivity with CBF and VTA DA neurons, we propose that the ventral striatum acts as a hub for integrating and computing prediction errors, conveying information about expected value to both neuromodulatory systems(*44, 45*). While here we emphasize the role of CBF prediction errors in learning, the CBF also supports cognitive performance via its effects on arousal and attention(*19, 46*). The governing principle of a reinforcement prediction error helps resolve how a single system could mediate such diverse functions, by orienting circuit preparation and adjustment towards reinforcement.

## Methods

### Animals

Adult C57BL/6 (JAX:000664), ChAT-IRES-cre (JAX:00641), and DAT-cre (JAX:006660) (over 2 months old, females and males) were used in accordance of the protocol approved by Cold Spring Harbor Laboratory Institutional Animal Care and Use Committee and National Institutes of Health guidelines.

### Viral injection and fiber implantation

Mice were anaesthetized with isoflurane (1-2%), head-shaved, and placed in a stereotax with the skull level (David Kopf Instruments). Thereafter, the eyes were protected with ophthalmic lubricant (Paralube Vet Ointment, Dechra Pharmaceuticals), lidocaine was subcutaneously injected under the scalp, and the skin cleaned and sterilized with a betadine solution. After cutting away the scalp, injection coordinates were marked and the skull was scraped and roughened using a bone scraper (Fine Science Tools), and craniotomies opened up using a dental drill. Coordinates were as follows: HDB 1.1 antero-posterior (AP) 0.8 medio-lateral (ML) −4.7 dorso-ventral (DV), VTA −3AP 0.8ML −4.5DV, BLA −1.2AP 3ML −4.2DV. AAV virus (300nL volume) was then pressure injected 100nL/min via a glass pipette pulled (P-97 Sutter Instruments) from borosilicate capillaries (Drummond calibrated 5ul; tip diameter, 20-50um). After a 10 minute period to allow the virus to diffuse, the pipette was slowly withdrawn and a fiber implant (400um core, 0.48NA, 2.5mm zirconia ferrule, 8mm fiber length, Doric Lenses) was mounted in a stereotaxic adapter, and lowered to the same depth as for viral injection. Then, the fiber was lightly secured with a UV adhesive (EM ESPE) such that the adapter could be removed and the skull and implant permanently bonded by successive coats of adhesive cement (C&V Metabond, Parkell). A titanium head bar was then placed anterior to the fiber implants and cemented in place with dental acrylic (Lang Dental). Following surgery, post-operative analgesia was maintained via intraperitoneal injections of buprenorphine (0.1mg/kg) for 48 hours. Mice were housed for a post-operative period of 3 weeks to allow for recovery and viral expression prior to the initiation of behavioral training and recording.

For initial HDB recordings, ChAT-cre mice were injected with AAV2.5 Syn-Flex-GCaMP6f (Addgene 100833). For dual fiber, reversal learning experiments, ChAT-cre/DAT-cre trans-heterozygotes were injected with AAV2.5 Syn-Flex-GCaMP6s (Addgene 100843) in the left HDB and either AAV2.5 Syn-Flex-NES-jRGECO1a (Addgene 100853) or later, to achieve improved signal to noise ratios, AAV2.1 Syn-Flex-NES-jRCaMP1a (Addgene 100848) in the right VTA. Importantly, we utilized contralateral fiber implants and red-shifted genetically-encoded calcium indicators for the VTA in order to exclude the possibility of picking up spurious signals from VTA axons projecting to the ventral striatum. For measurement of extracellular acetylcholine, wild type mice were bilaterally injected with AAV ACh3.0, an improved variant of a recently described fluorescent acetylcholine sensor(*26*).

### Behavior

Mice were water-restricted prior to training to achieve 85%–90% of body weight. To habituate mice to head-fixation, un-cued water reward (5 mL) was delivered at randomized intervals (exponential, mean = 3 s) for 1 or 2 sessions and licking behavior monitored using a custom lickometer with an IR beam-break to register lick events. Mice were trained in head-fixed, Pavlovian, odor-cued tasks, i.e. reinforcement outcomes were deterministic after trial initiation. Prior to trial initiation, odor and reinforcement stimuli were selected pseudorandomally from probabilistic contingency tables. Following habituation, mice progressed to a simplified training stage in which a single odorant predicted water reward (8uL unless otherwise specified) with 90% probability, and with a 1s trace delay separating odor and outcome. Typically after 1 or 2 training sessions, mice developed robust anticipatory licking in response to odor delivery, at which point they graduated to other task variants. In 2 odor valence tasks, a second odor was introduced that was associated with an aversive stimulus, either a mild air puff to the eye or tail shock. For the high/low value task, the high and low value odors were associated with reward/air puff/omission probabilities of 0.8/0.1/0.1 and 0.35/0.55/0.1, respectively.

Unless otherwise stated, each trial consisted of a 4s baseline period, a 1s odor stimulus, a 2s trace delay, and a 4s post-outcome period of photometry recording. Trials were separated by a randomized interval (truncated exponential, mean=6 s, minimum=2s, maximum=18s). Odors were diluted in mineral oil and delivered at a final concentration of 1% using a custom olfactometer controlled by an Arduino Uno microprocessor. Monomolecular odorants were randomly assigned to different reinforcement contingencies across behavioral tasks and included isoamyl acetate, ethyl tiglate, cineole, and 1-hexanol.

Two reversal learning tasks were employed involving 1) valence reversals and 2) value reversals. In the valence task, the “CS+” odor was positively valued, i.e. overall predictive of reward with reward probability set at 0.9. Thus, the CS+ odor drove conditioned behavioral responses-anticipatory licking-that were used to index learning. In contrast, the CS-odor was negatively valued (opposing valence), being predictive of punishment with a probability set to 0.5. In the valence task, odor contingency reversals were manually determined by the experimenter, occurring after ∼100 trials in each session. In the value task, reward occurred more frequently for the CS+ odor (p=0.8) compared to the CS-(p=0.05). Furthermore, reversals were controlled automatically according to a runny tally (computed every trial) of whether the anticipatory lick rates of the last 10 CS+ and the last 10 CS-trials since the previous reversal were statistically discriminable. When 90% of the P values over the last 20 trials (all trial types) exceeded significance, a reversal was triggered. P values for this tally were computed by comparing the auROC value (area under the receiver operator characteristic curve) between CS+ and CS-trials to a null distribution of auROC values generated by repeatedly shuffling trial labels. Furthermore, a 50 trial minimum number of post-reversal trials and a minimum response probability of CS+ trials of 0.7 across the last 20 trials was further imposed.

To compare externally and internally driven neuromodulator dynamics, a random rewards task was utilized in which occasional water rewards (8uL) were delivered with inter-reward intervals drawn from a truncated exponential distribution (30s mean, 2s minimum, 120s maximum). Mice were placed on a platter-type wheel coupled to a rotary encoder to measure running velocity. Furthermore, separate video cameras (Point Grey) were used to record pupil dilation and whisking. An infrared light was used to provide oblique illumination to the eye. A blue LED was placed opposite the animals head and adjusted to produce background light to the opposite eye sufficient to induce moderate pupillary contraction. Whisking movement was quantified as the square of the difference between pixel values between successive image frames. Pupil diameter was calculated using using a custom Matlab program. After choosing a manually selected threshold to isolate the pupil, morphological dilation/erosion was used to consolidate the pupil region after which linear regression was used to calculate the diameter of the best fit circle to the perimeter of the pupil region.

### Fiber Photometry

For fiber photometry, a 490nm LED light source (M470F3 Thorlabs) was collimated via an aspheric condenser (ACL25416U Thorlabs), passed through an excitation filter (ET470/24M Chroma), bounced off a dichroic mirror (T495LPXR Chroma), and launched into a 400mm core, 0.48NA fiber patch cable using an aspheric objective lens (A240TM-A Thorlabs). Fluorescence excitation and detection were both accomplished through one multimode optical fiber. Then fluorescence was passed through an emission filter (ET525/50M Chroma), focused with a plano-convex lens (f = 30 LA1805-A Thorlabs), and collected using an amplified photodiode (IM 2151 New Focus). The fluorescence signal was amplitude-modulated by sinusoidally varying the command voltage of the LED driver (LEDD1B Thorlabs) and decoded in silico(*47*). Data were acquired using a data acquisition card (PCIe-6321 National Instruments) and analyzed using custom MATLAB code. Olfactometer, photometry data acquisition, and behavioral monitoring were controlled and synchronized using a flexible, open-source master controller interfacing with MATLAB (Bpod State Machine, Sanworks LLC).

### Analysis

Fluorescence signals were expressed in units roughly equivalent to meausurement SNR as Fσ_bl_, where Fσ_bl_=(F-F_bl_)/σ_b□_, and F_bl_ is the mean fluorescence value for a given trial across the 4s baseline acquisition period and σ_bl_ is the baseline period standard deviation averaged across all trials within a given session. Prior to any analysis being performed, each session was manually truncated to exclude trials where the mouse had reached a state of satiety by plotting the number of reward licks vs. rewarded trials and selecting a trial before the lick rate steeply declined. Fluorescence responses to odor stimuli and reinforcement were estimated as mean values across 1s windows following stimulus delivery. Owing to the prolonged decay of cue-evoked fluorescence responses observed using GCamP6f in the HDB, reward responses were baselined using a 0.5s period immediately preceding reward delivery for the quantification of surprise modulation in the context of the high/low value probabilistic outcome task (Fig. 1J). For reversal experiments, optimized cue window parameters (window start and stop) were determined by grid search in order to maximize the discriminability of CS+ and CS-odor-evoked responses. Reversals were included for analysis according to the following criteria: 1) CBF and VTA DA cue responses to the new CS+ odor increased following the reversal. 2) Conditioned licking discriminated between CS+ and CS-odors following the reversal (same criterion for automated triggering of value reversals, see above). Changepoint trials corresponded to the maximum vertical difference between the cumulative sum of cue-evoked responses (starting from 20 trials prior to the reversal point) and a line through the origin that coterminated with this sum.

### Retrograde Viral Tracing

For transsynaptic retrograde tracing, ChAT(cre/+); DAT(cre/+) mice were injected intracranially with 300nL of helper virus cocktail, comprised of a 2:1 ratio of Rabies Glycoprotein to TVA (AAV2/9-CAG-Flex-mKate-T2A-N2c-G, titer 1E12 VP/mL; AAV2/9-CAG-Flex-mKate-T2A-TVA, titer 1E12 VP/mL) injected unilaterally at both the basal forebrain (coordinates in mm relative to bregma, AP −0.1, ML +1.3, DV −5.0) as well as at the ventral tegmental area (coordinates in mm relative to bregma, AP −3.0, ML +0.8, DV −4.3). After three weeks, 500nL of pseudotyped rabies was injected unilaterally at both the basal forebrain and the ventral tegmental area (EnvA-Gdeleted-Rabies-EGFP, titer 1E9 VP/mL;EnvA-Gdeleted-Rabies-mCherry, titer 1E9 VP/mL;).

