## Supplemental for "Basal forebrain-derived acetylcholine encodes valence-free reinforcement prediction error"

**Figure S1**

CBF responses to reward-predictive cues are acquired with learning

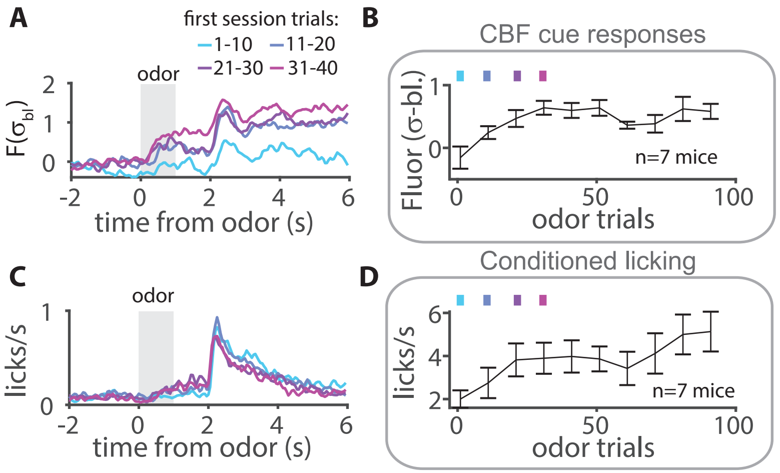


**A** Average CBF responses during initial training for example mouse. **B** Cue responses are initially absent but strengthen with training (n=7 mice). **C** Increase in anticipatory licking during initial conditioning for example mouse. **D** Summary data shows that lick responses are acquired with learning (n=7 mice).

**Figure S2**

Value-related CBF activity during cue period is most strongly associated with stimulus, not lick onset

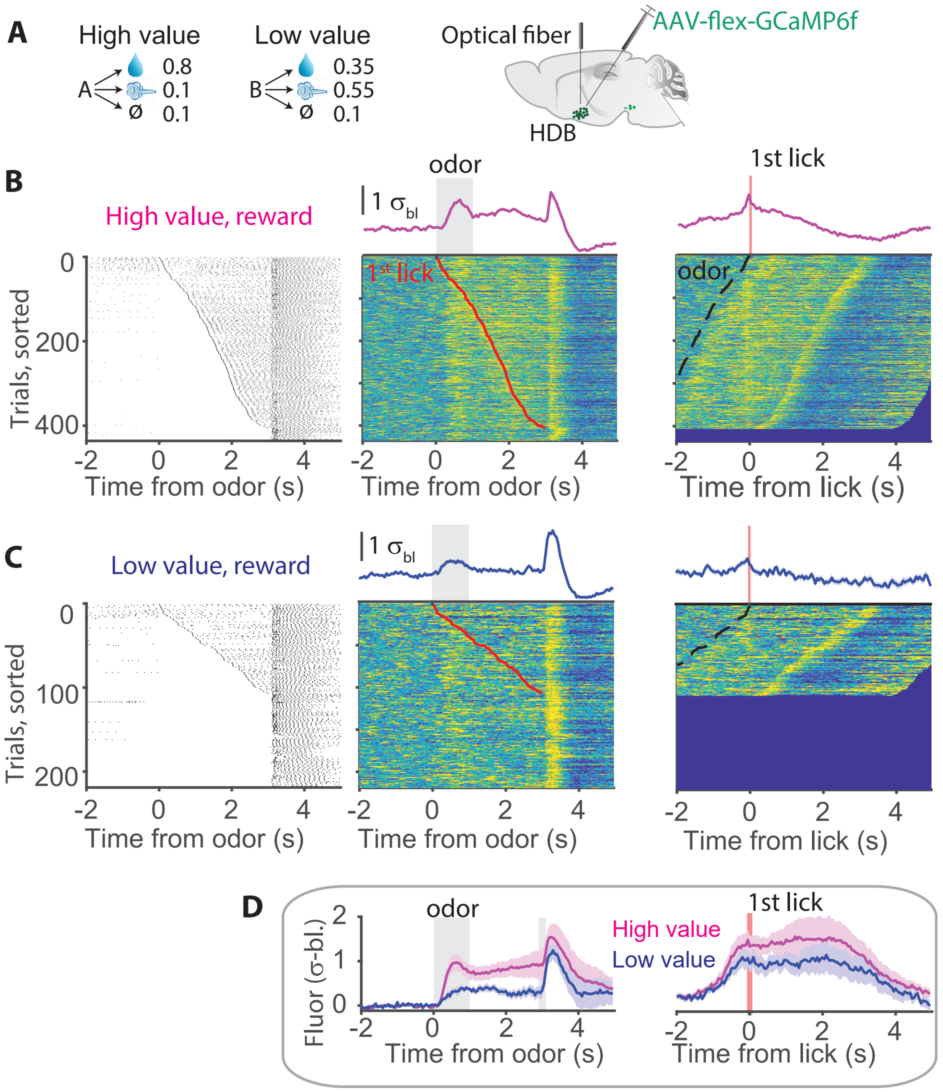


**A** Schematic for probabilstic outcome task and photometry recordings of CBF activity. **B** Data for high value, reward trials for example mouse. Left, lick raster sorted by latency to first lick during the odor cue and trace periods. Middle, photometry raster aligned to odor onset with average response depicted above and lick onset denoted by red line. Right, photometry raster aligned to first lick onset and corresponding average. **C** Same as for B but for low value, reward trials. **D** Summary data comparing alignment to odor vs lick onset, n=7 mice.

**Figure S3**

Consistency of reinforcement-driven extracellular ACh release

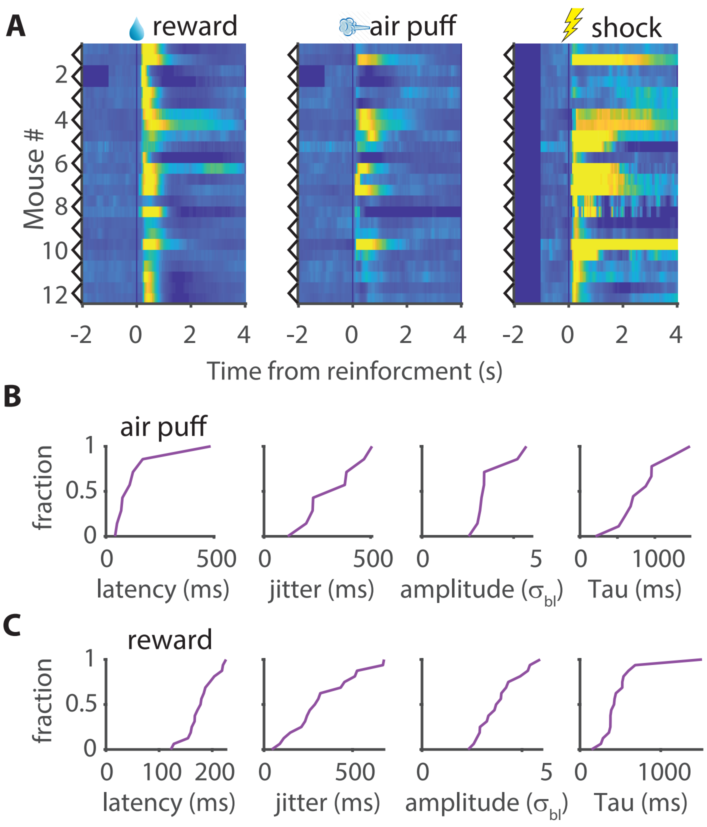


**A** Average fluorescence responses of BLA principal neurons expressing the acetylcholine sensor, ACh4.3 to reward, air puff, and tail shock delivery (n=12/24 mice, recording sites). **B** Cumulative histograms quantifying response latency, jitter, amplitude and decay time constant for air puff trials. **C** Same as for B but for reward delivery.

**Figure S4**

Cue responses of CBF and VTA DA neurons show similar dynamics during serial value reversals

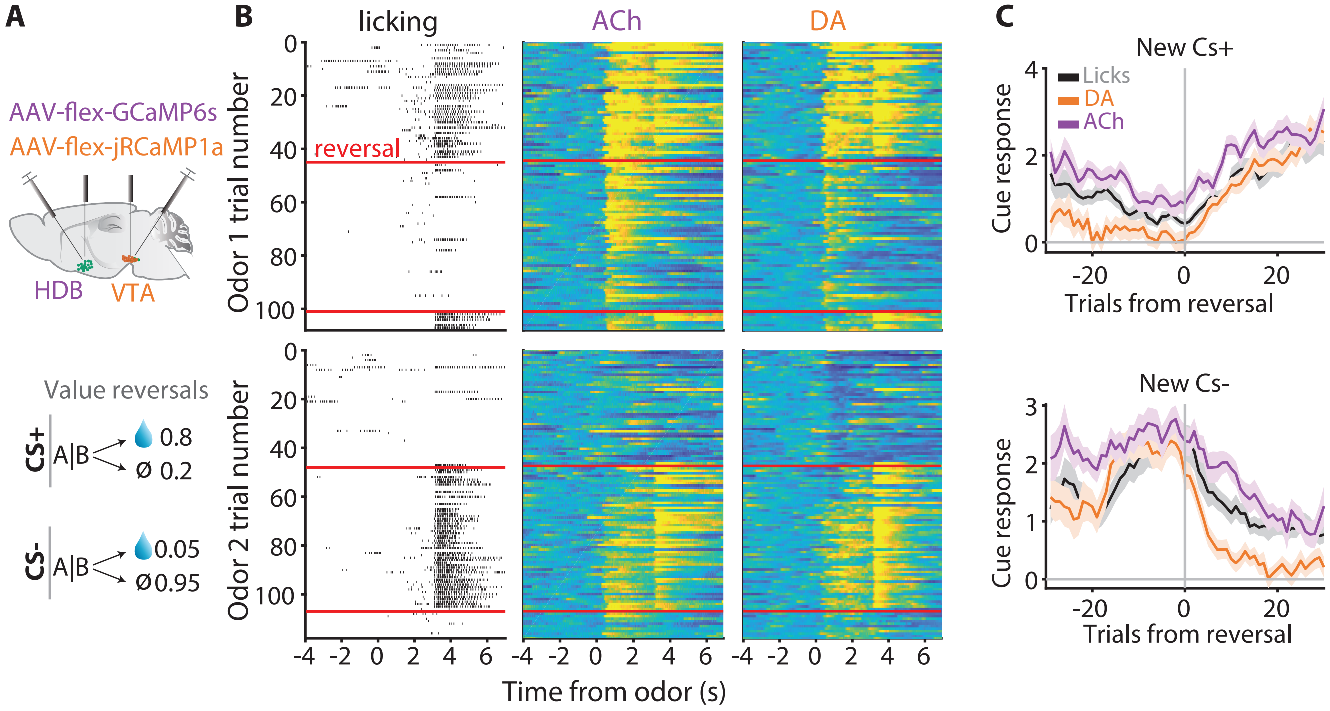


**A** Recording and task schematic for serial value reversals. **B** Example raster plots for licking (left), CBF (middle) and VTA DA (right) photometry, shown separately for odor 1 (top) and odor 2 (bottom) trials. Red lines denote reversal points. **C** Average cue-evoked licking, CBF and VTA DA measurement relative to reversal trial (n=16/21 New CS+/New CS- reversals).

**Figure S5**

Behavioral changepoint detection and alignment of neural data

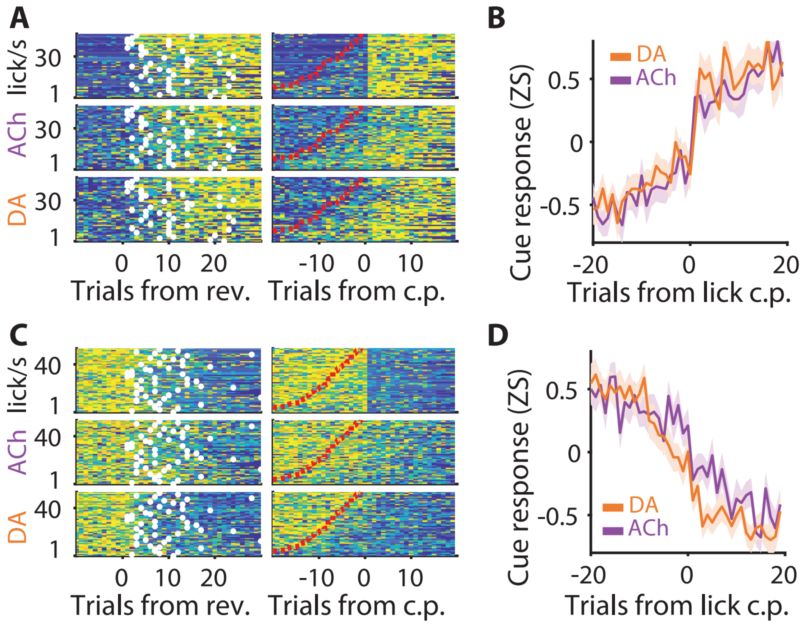


**A** Raster images across reversals quantifying anticapatory licking (top), CBF (middle), and VTA DA (bottom) activity in response to delivery of new CS+ odor. Images at left are aligned to reversals with changepoints computed from lick rates indicated by white markers. Images at right are aligned to changepoints and sorted by learning latency with preceding reversal indicated by red dotted line. **B** Indistinguishable average time courses for acquisition of CBF and VTA DA responses to new CS+ odor aligned to lick changepoint. **C** As for A but for new CS- trials (extinction). **D** VTA DA responses decline more rapidly relative CBF responses during extinction learning.
